# Factors Influencing Monoclonal Antibody Pharmacokinetics Across Varying Immune Perturbations

**DOI:** 10.1101/2025.04.29.651349

**Authors:** Jonathon DeBonis, Anthony Davis, Zeshi Wang, Cody Fell, Michael Diehl, Oleg A. Igoshin, Omid Veiseh

**Author notes:** Corresponding Authors: Oleg Igoshin, Department of Bioengineering, Rice University, Houston, Texas, USA & Omid Veiseh, Department of Bioengineering, Rice University, Houston, Texas, USA.

## Abstract

The development of continuous-release devices or injectables for the long-term delivery of biologics is of great interest, especially monoclonal antibodies (mAbs) that require frequent, high-dose injections. Preclinical testing of these technologies in murine models is necessary for clinical translation; however, xenogeneic responses to the mAb and foreign body responses to the implants or injectables can confound results. Immune system knockout (KO) models that affect immune cells are often used in these experiments, but the effects of KO models on mAb pharmacokinetics (PK) are not well characterized. Here, we investigated the PK profile of the human mAb 3BNC117 after intravenous, subcutaneous, and intraperitoneal injections in four mouse strains: BL6, BCD, RAG2, and NSG mice. Noncompartmental analysis was used to quantify differences in PK between each mouse strain. Strikingly, both BL6 and NSG mice exhibited significantly higher mAb clearance compared to the other two strains. To better understand these differences, we developed a minimally physiological based PK model of mAb PK and estimated model parameters using nonlinear mixed effects modeling. The fitted model parameters illustrated how specific processes change in each strain, including the change in clearance rates over time in BL6 and differences in mAb lymphatic uptake in NSG mice. We then used simple allometric scaling relationships to assess which strains were reasonably predictive of human mAb PK. NSG and BL6 mice were found to be unpredictive of human PK, unlike RAG2 and BCD mice. Overall, these results highlight the importance of selecting an appropriate KO strain for preclinical mAb evaluation and demonstrate that RAG2 and BCD strains are suitable mouse models for investigation of mAb PK.

**Graphical Abstract:** 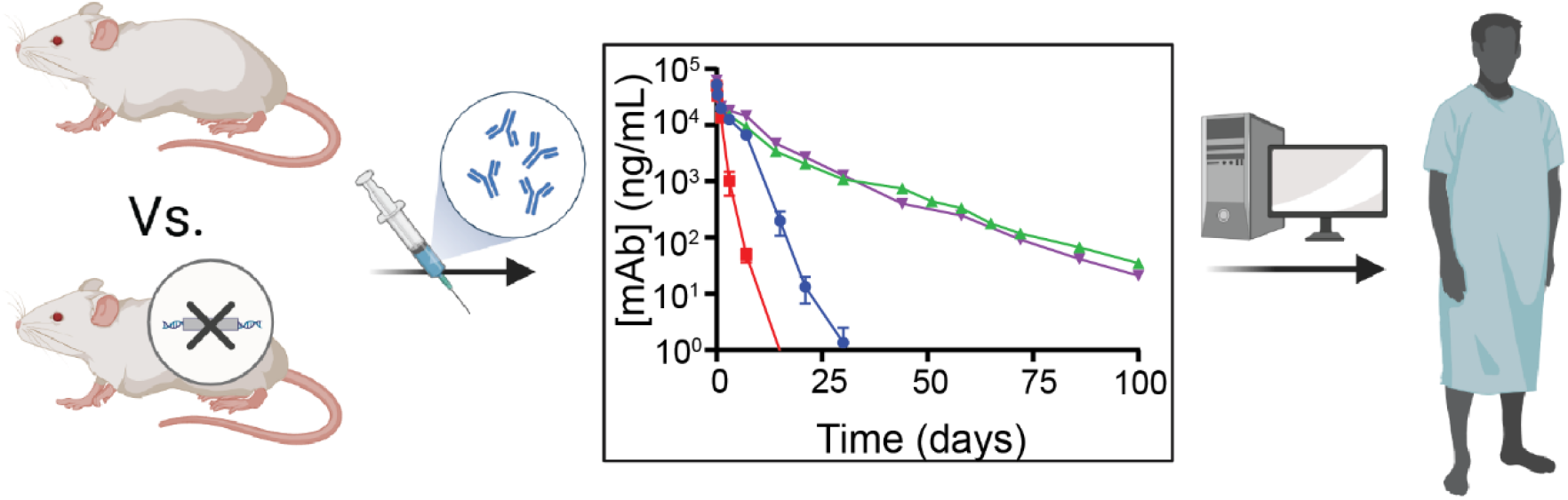

## Introduction

Monoclonal antibodies (mAbs) are a highly versatile biotherapeutic modality with the potential to treat many different patient indications. In 2023, 20% of newly approved drugs were mAbs, primarily due to their high target specificity, diverse range of functionality, and serum stability [1,2]. Engineering of the mAb variable region (Fab) allows for a broad range of target proteins to be selected, and the effect of target binding can be manipulated by engineering of the constant region (Fc). Thus, mAbs are capable of treating a wide range of diseases [3,4]. There are many approved mAbs for the treatment of cancer and autoimmune disease [5], and others are in clinical development for the treatment of other indications such as HIV [6], Malaria [7], COVID-19 [8], bacterial sepsis [9], and even Alzheimer’s disease [10]. However, mAbs are typically administered as intravenous (i.v.) or subcutaneous (s.c.) injections, requiring large and frequent doses for long-term efficacy. For this reason, technologies for the sustained and controlled release of mAbs are of great interest due to their ability to maintain more durable and consistent therapeutic concentrations over the course of treatment [4,11,12]. These technologies often take the form of implantable devices [13–15] or injectable hydrogels [16–18] that, when injected into peripheral tissue, continuously release mAb such that exposure is maintained for extended periods of time.

Any new continuous-release platform needs to be tested in appropriate preclinical models to verify the durability of the technology and pharmacokinetic (PK) profile of the released mAb. Murine models are a vital step in this characterization, especially for implantable devices, where it is difficult to predict how an implant and its released substance will perform in the complex *in vivo* environment. For example, host responses such as the foreign body response (FBR) against the implant and the anti-drug antibody (ADA) response against the released substance can make interpreting experimental data difficult [19–21]. Several murine models are often needed to systematically isolate facets of implanted device performance. For example, if one wishes to test the fibrotic axis of FBR on the profile of released mAb in the absence of ADA response, a mouse model such as the B6.129S2-Ighmtm1Cgn/J strain, which lacks functional B cells, would prevent the development of an ADA response [22]. In other studies, such as those evaluating implant biocompatibility, a general-purpose healthy model such as the C57BL6 strain is useful [23]. Utilizing murine models with varying levels of immunodeficiency caused by genetic perturbations, often by gene knockout (KO models), is a valuable method to characterize continuous-release device performance. However, caution is needed to avoid making incorrect assumptions when only one KO model is used.

Immune system KO can affect mAb PK. For example, in fully immunocompetent mice such as the C57BL6 strain, human mAbs elicit an ADA response, which leads to a rapid increase in mAb clearance and makes the long-term study of mAb PK difficult. KO models are known to mitigate the formation of ADA, prolonging treatment duration [24]. However, severely immunocompromised (e.g., NSG) mice exhibit increased mAb clearance and abnormal biodistribution following bolus i.v. injection [25,26]. This phenomenon appears to occur in a FcγR-dependent manner, but the impact on e.v. administration has not been characterized.

Here, we investigate the effects of immune system KO on mAb PK following bolus i.v., s.c. and intraperitoneal (i.p.) injections. To this end, we evaluated the PK of 3BNC117 following bolus i.v., s.c., and i.p. injections in four mouse strains with varying levels of immune KO. 3BNC117 is a fully human mAb that binds to the GP120 spike protein of HIV and has been shown in clinical trials to neutralize HIV in infected patients [27,28]. A minimal physiologically based PK (mPBPK) model incorporating mouse weight and strain as covariates is used to quantify how specific absorption, distribution, and elimination properties of 3BNC117 are influenced by immune KO. The model is then combined with simple allometric scaling to assess which mouse strains exhibit mAb PK relevant for clinical translation.

## Methods

### Reagents

The purified 3BNC117 mAb protein used in the in vivo studies described was provided by the Duke Human Vaccine Institute Protein Production Facility and facilitated by the Collaboration for AIDS Vaccine Discovery and the Gates Foundation. This mAb was originally isolated from an HIV-infected donor and shown to broadly neutralize across HIV clades with high affinity [29]. As such, it is a fully human IgG1 mAb, whose sequence was determined and used to produce recombinant mAb for this study. The total Human IgG enzyme-linked immunosorbent assay (ELISA) kit used for the quantification of 3BNC117 serum levels was purchased from Abcam (Cat No: ab195215). All other reagents and consumables used in the study were obtained from various commercial sources.

### Animals

C57BL6/N (BL6) mice (8 weeks old, female) were purchased from Charles River Laboratories. B6.129S2-Ighmtm1Cgn/J (BCD) mice, B6.Cg-Rag2tm1.1Cgn/J (RAG2) mice, and NOD.CgPrkdcscid Il2rgtm1Wjl/SzJ (NSG) mice (8 weeks old, female) were purchased from Jackson Laboratories.

### Animal Experiments

Animal experiments in this study were performed in accordance with the Guidelines for the Care and Use of Laboratory Animals at Rice University, which is accredited by the Association for Assessment and Accreditation of Laboratory Animal Care (AAALAC) International.

### Evaluation of 3BNC117 PK in Mice

The PK of 3BNC117 in murine serum was evaluated by administering bolus i.v., s.c., and i.p. injections at a fixed dose of 56 μg. The drug solution consisted of purified 3BNC117 antibody in sterile Dulbecco’s phosphate-buffered saline and was administered at a volume of 100 μL. Animals were excluded from subsequent blood collection and analysis if i.v., s.c., or i.p. administration was incorrectly performed, leading to some variability in sample size between experimental groups. Blood was collected from the saphenous vein without anesthesia. Blood samples were taken at 1 hour, 4 hours, 1 day, 3 days, 7 days, and ∼weekly thereafter until blood concentrations fell below the accuracy threshold of the ELISA (2.3 ng/mL due to dilution) or 100 days was reached. Blood samples were allowed to coagulate for 1 hour at room temperature, then centrifuged at 3,500xG for 10 minutes to allow the collection of serum. The collected serum was stored at -80 C until analysis via ELISA.

### Enzyme-linked immunosorbent assay

Total Human IgG ELISA kits obtained from Abcam were utilized, and the manufacturer’s protocol was followed. Samples were diluted into the dynamic range of the assay with the supplied sample diluent. Following the addition of the sample to supplied 96-well plates, the addition of capture and detector antibody, washing, 15-minute incubation with tetramethylbenzidine (TMB), and addition of the stop solution, the absorbance was measured at OD450 nm using an infinite M200 Pro plate reader (Tecan). Concentrations were then calculated using standard curves.

### Noncompartmental Analysis

Noncompartmental analysis (NCA) was performed using the ‘NonCompart’ and ‘ncar’ packages in R [30]. Maximum observed concentration (C_max_), the area under the concentration v. time curve (AUC), clearance (CL), terminal half-life (t_1/2_), absolute bioavailability (F) and steady-state volume of distribution (V_SS_) were calculated for each mouse. Values for each NCA parameter were compared based on mouse strain.

### Statistical Analysis

NCA parameters were compared across experimental groups. Data was plotted as mean ± standard deviation in bar charts. Statistical differences between groups were assessed using two-way ANOVA for those data sets comparing administration routes and strain, while one-way ANOVA was utilized for strain differences alone. Each mouse strain was compared to the BL6 strain by post hoc multiple comparisons tests using Dunnett’s test within administration routes if applicable. Statistical analysis for outlier exclusion was performed on data points that were visually aberrant using ROUT outlier detection with Q = 1% within administration routes, treating each time point as a group. These statistical analyses were performed using GraphPad Prism software (version 10.3.1, for Windows, GraphPad Software, Boston, Massachusetts USA, www.graphpad.com). Statistical significance was defined as *p* < 0.05.

### Minimal Physiologically Based Pharmacokinetic Model of Bolus mAb Injection

An mPBPK model of bolus mAb injection, absorption, distribution, and elimination was developed based on several models in the literature [31–34]. The model consists of six total compartments: the central/blood compartment (B), the peripheral tissue compartment (T), the peripheral lymphatic compartment (P), the central lymphatic compartment (L), the subcutaneous dosing compartment (S), and the intraperitoneal dosing compartment (I). The concentration of mAb in each compartment is described by the following ordinary differential equations:

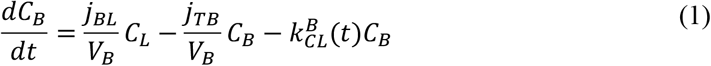

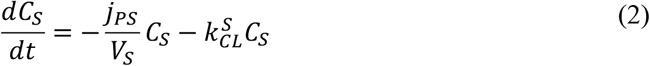

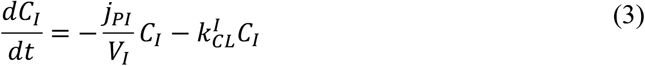

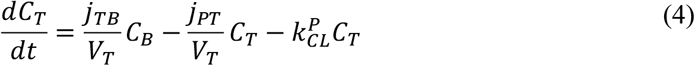

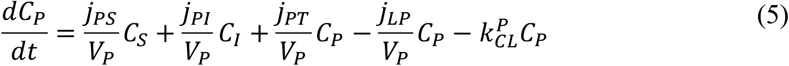

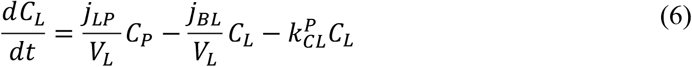

Here, *C*_*x*_ is the concentration of mAb in compartment *x* (ng/mL), *V*_*x*_ is the volume of compartment *x, j*_*yx*_ is the rate of mAb-carrying fluid flow from compartment *x* to compartment *y*, and 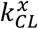 is the first order clearance rate constant from compartment *x*. Bolus injection of mAb was represented by setting the initial concentration of mAb in the corresponding compartment according to:

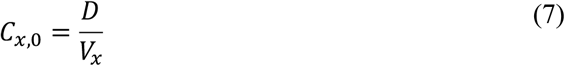

where *D* is the dose administered (ng). Mouse strain and starting weight were included in our model as covariates. Weight influenced volumes and all elimination rate constants according to:

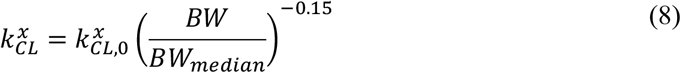

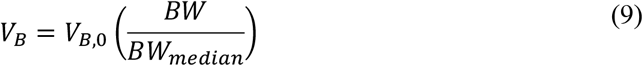

where *BW* is the mouse weight and *BW*_*median*_ is the median weight of all mice. Mouse strain influenced the time-dependence of clearance from the central compartment and the efficiency of mAb reabsorption into the blood following distribution into peripheral tissue. Time-dependent clearance was represented based on other models in the literature [32,33] according to:

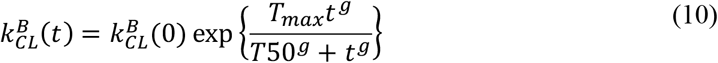

Where *T*_*max*_ determines the maximum log fold change, *T*50 is the time at which clearance reaches 50% of its maximum log fold change, *g* is a hill coefficient, and 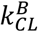 (0) is the value of the clearance rate constant at time zero. BL6 mice were given a time-dependent increase in elimination (positive *T*_*max*_), and RAG2 and BCD mice were given a time-dependent decrease in elimination (negative *T*_*max*_). Impaired ability of NSG mice to reabsorb mAb from peripheral tissue is represented by a scaling of the fluid flow rate *j*_*PT*_ by the factor *δ*_*NSG*_, such that *j*_*PT,NSG*_ = *δ*_*NSG*_*j*_*PT*_.

### Parameter Estimation with Nonlinear Mixed Effects Modeling

In total, the developed model has 23 parameters (**Tables S2**), with 7 fixed parameters based on literature values or best approximations and 16 parameters estimated via fitting to data. For model fitting, measured values below the limit of quantification (LOQ) but above the limit of detection (LOD) were included for parameter estimation as described in the literature [35]. Model parameters and inter-individual variability (IIV) were estimated using nonlinear mixed effects modeling with the nlmixr package in R [36–38] using the FOCEI estimation method, log-normally distributed parameters, and both proportional and additive error. Random effect parameters were selected by first estimating model parameter with only random effect parameters, and down-selecting based on calculated shrinkage values and analysis of Emprical Bayes Estimates (EBEs) for the presence of strain-dependence.

## Results

### Immune System Perturbations Significantly Alter Monoclonal Antibody Pharmacokinetics

To interrogate the effect of immune KO on 3BNC117 PK, purified 3BNC117 was administered to four strains of mice: C57BL6 (BL6), B6.129S2-Ighmtm1Cgn/J (BCD), B6.Cg-Rag2tm1.1Cgn/J (RAG2), and NOD.CgPrkdcscid Il2rgtm1Wjl/SzJ (NSG). BL6 mice are a fully immune competent general purpose mouse model, while the other strains contain targeted contain genetic mutations that cause immune cell perturbations: BCD mice lack mature B cells, RAG2 mice lack mature T and B cells, and NSG mice are extremely immunodeficient, lacking functional macrophages, dendritic cells, hemolytic complement system, T cells, B cells and natural killer cells (**Figure 1A**). Doses were administered as bolus i.v., s.c., and i.p. injections, and the serum concentration of 3BNC117 was tracked over time (**Figure 1B-D**). Serum was collected at 1 hour, 4 hours, 24 hours, 3 days, 7 days, and approximately weekly thereafter until the study date completion or until the concentration fell below the LOQ of the assay. The concentration of mAb in serum samples was determined using a human IgG ELISA (Abcam 195215). Comparison of the 3BNC117 concentration vs. time curves following 3BNC117 dosing in each mouse demonstrates that immune KO has varying but significant effects on 3BNC117 PK. In BL6 mice, 3BNC117 follows a typical mAb PK profile consisting of rapid distribution and slow elimination phases [39,40] until roughly one week, when mAb elimination rapidly increases. 3BNC117 PK in RAG2 and BCD mice, however, does not exhibit this rapid increase in elimination and is similar in both strains. In stark contrast to all other strains, NSG mice exhibited a rapid decrease in serum mAb concentrations immediately after dosing and fell below the LOQ of the assay roughly one week after dosing.

**Figure 1:**
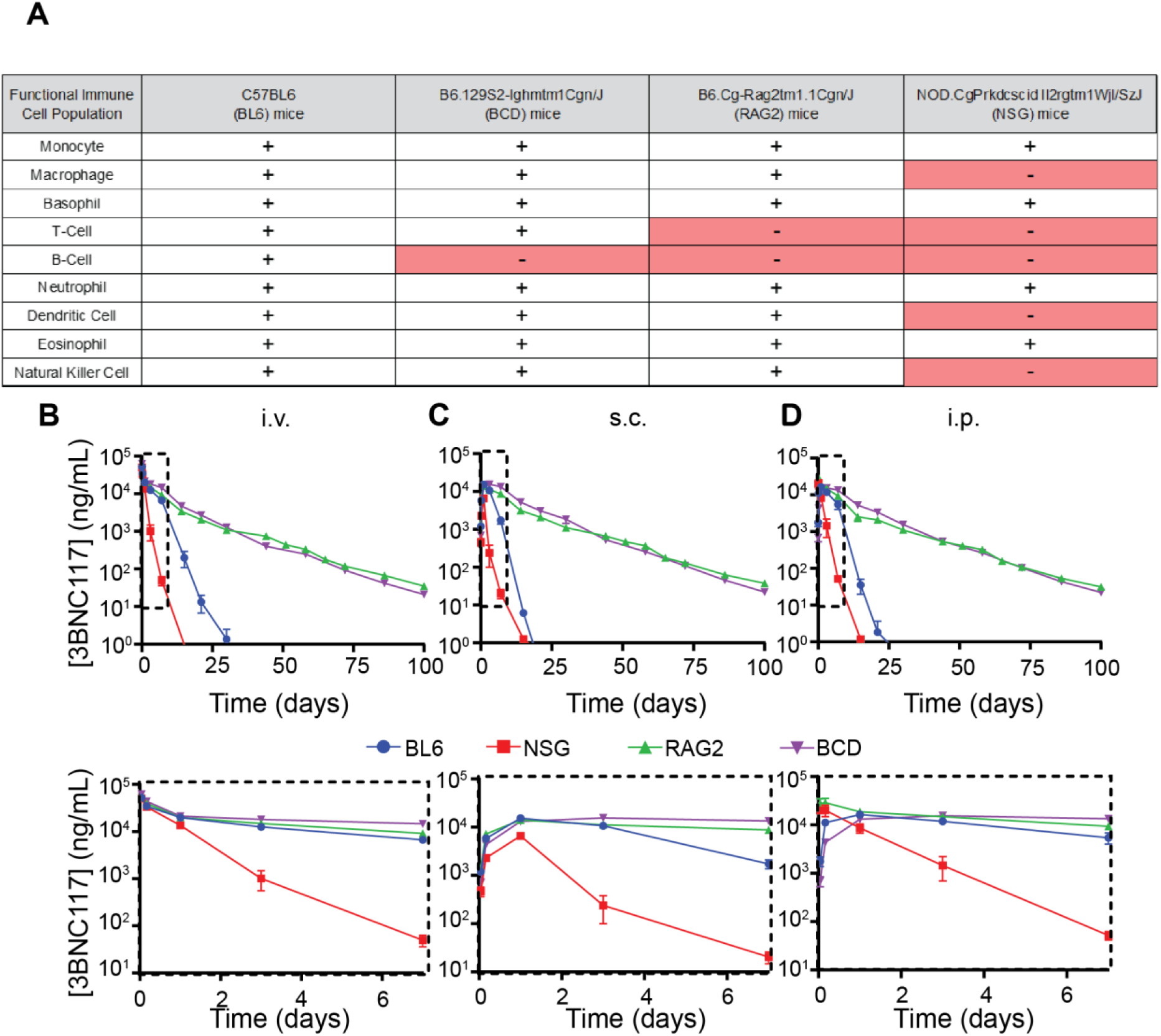
Concentration v. time profiles following bolus injection of 56 µg purified 3BNC117. A) Matrix summarizing the effects of immune KO in each mouse strain. B-D) Serum 3BNC117 concentration vs time data collected from all strains of mice following bolus B) i.v. C) s.c. or D) i.p. injections. Blue points: BL6. Red points: NSG. Green points: RAG2. Purple points: BCD.

To quantify the observed differences in mAb PK between strains, we calculated key NCA parameters associated with each mouse strain, including *C*_*max*_, *AUC*, bioavailability (*F*), *CL*, and *t*_*1/2*_ (**Supplementary Table S1**). Following i.v. administration, *C*_*max*_ values for all four mouse strains were similar, but s.c. and i.p. administration led to significantly different values across strains. Furthermore, *C*_*max*_ following i.p. administration was higher than after s.c. administration in all cases (**Figure 2A**). Despite similar *C*_*max*_ values, calculated *AUC* values varied across mouse strains, with NSG mice exhibiting significantly reduced values compared to BL6 mice. RAG2 and BCD mice exhibited significantly increased AUC compared to other strains (**Figure 2B**). Similarly, bioavailability (*F*) values varied across mouse strains and were improved following i.p. injections compared to s.c. injections. (**Figure 2C**). NSG and BL6 mice also exhibited much higher clearance rates and shorter half-lives than RAG2 and BCD mice (**Figure 2D-E**).

**Figure 2:**
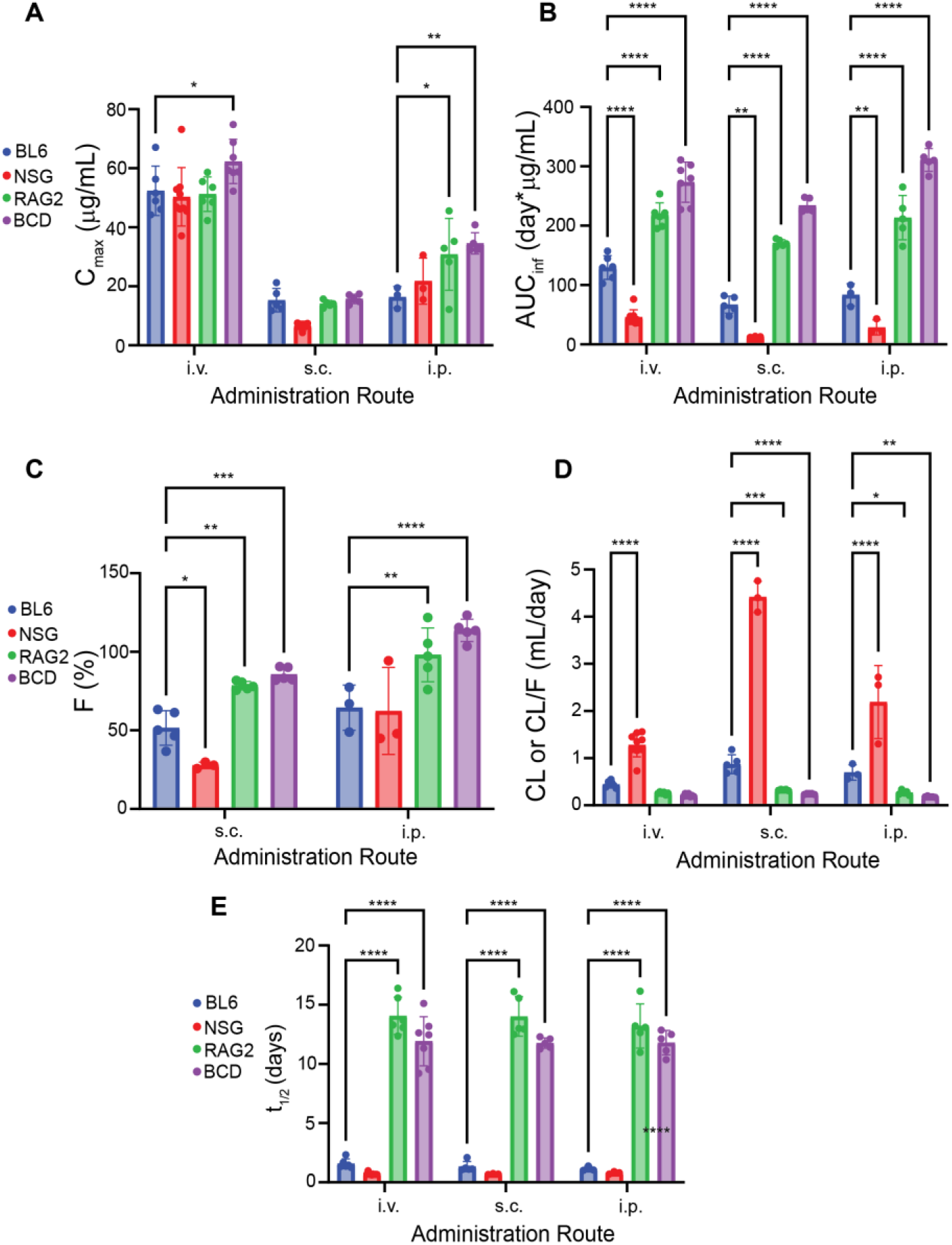
Calculated NCA parameters following i.v., s.c., and i.p. administration in each mouse strain. A) Peak measured concentration (C_max_), B) Area under the curve (AUC), C) Bioavailability (F), D) Clearance (CL or CL/F), E) Half-life.

We hypothesize that the differences in the PK profiles across these mouse strains were primarily driven by two effects: ADA formation in BL6 mice and impaired reabsorption of mAb to the blood in NSG mice. ADA formation in BL6 mice is known to occur following dosing of human mAbs [24]. In NSG mice, our hypothesis is based on the findings of Sharma et al. [25] and Li et al. [26], where it was demonstrated that mAbs in NSG mice tend to accumulate in peripheral tissues in an FcγR-dependent manner. Specifically, we hypothesize that significant binding to FcγR in peripheral tissues during the rapid distribution phase of mAb PK leads to the inability of these mice to reabsorb mAb via the lymphatic system, causing an apparent increase in clearance. There is also evidence in the literature demonstrating that the lymphatic system architecture in NSG mice is reduced in size compared to other mouse strains, which may also influence the ability to reabsorb mAbs [41].

### PK Data is Accurately Represented by a Minimal Physiologically Based PK Model

To determine if our proposed mechanisms can account for the trends observed in data and to quantify the potential effects of these mechanisms, we developed an mPBPK model representing mAb dosing via bolus i.v., s.c., or i.p. injection, distribution into peripheral tissues, lymphatic transport from peripheral tissues to circulation, and clearance from each compartment based on similar models in the literature [31–34] (**Figure 3, Equations 1-10**). ADA formation in BL6 mice was incorporated into our model empirically with a time-dependent increase in clearance from the central compartment. Furthermore, RAG2 and BCD time courses were best represented with a clearance rate that decreases at long times, but this mechanism is unclear. Impaired lymphatic reuptake of mAb in NSG mice is represented by the parameter *δ*_*NSG*_, the fold-decrease in lymph fluid uptake from the peripheral tissue compartment to the peripheral lymphatic in our model (*j*_*PT*_). Other than the scaling of this parameter, all model parameters, except those adding time-dependent clearance, were assumed to be identical in NSG mice to other strains.

**Figure 3:**
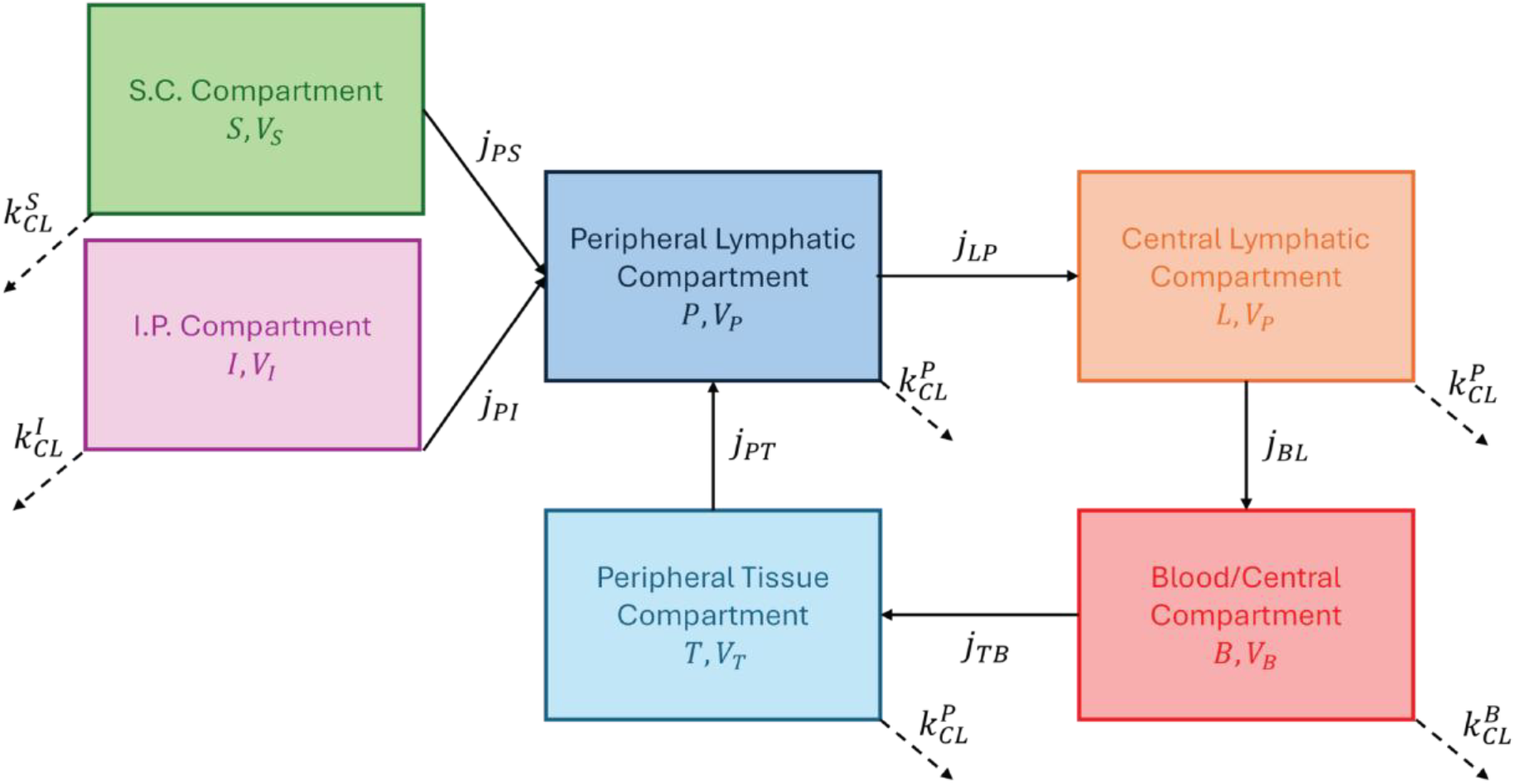
Schematic of mPBPK model developed to represent i.v., s.c., and i.p. mAb injection.

Model parameters were estimated using nonlinear mixed effects modeling with mouse strain and body weight as covariates. Initial parameter estimates were determined with the naïve pooled approach [42], and parameters were estimated using the FOCEI estimation method in the nlmixr package in R. Parameter estimation results in model predictions that are in good agreement with experimental data (**Figure 4, Supplementary Figure S1**). The best-performing model contained both mixed and random effect parameters, with 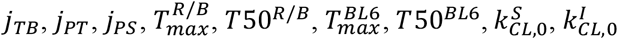 and 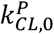 having inter-individual variability (IIV) (**Table 1, Supplementary Figure S2**). Specifically, the mechanisms incorporated in our model successfully capture the time-dependent increase observed in BL6 mice (**Figure 4A**) and the rapid initial decrease in serum mAb observed in NSG mice (**Figure 4B**). In the absence of these aspects, the PK profiles of 3BNC117 are well-captured in both RAG2 and BCD mice (**Figure 4C-D**).

**Table 1:** Estimated fixed and random effect parameters.

| Fixed Effect Parameters |  |  |
| --- | --- | --- |
| Parameter |  | Estimated Value |
| $\delta_{NSG}$ | | 6.1e-5 |
| $j_{PI}$ | | 80 /day |
| $k0_{CL}^B$ | | 0.15 /day |
| $g$ | | 8.2 |
| $g_{BL6}$ | | 3.5 |
| $V_{B,0}$ | | 1.25 mL |
| $\sigma_{prop}$ | | 0.29 |
| $\sigma_{add}$ | | 0.594 ng/mL |
| Random Effect Parameters |  |  |
| Parameter | Estimated Value | BSV* |
| $j_{TB}$ | 1.28 mL/day | 12 % |
| $j_{PT}$ | 0.050 mL/day | 28 |
| $j_{PS}$ | 0.15 mL/day | 46% |
| $T_{max}^{R/B}$ | -1.0 | 16% |
| $T50^{R/B}$ | 25 days | 18% |
| $T^{BL6}$ | 2.8 | 21% |
| $T50^{BL6}$ | 5.2 days | 15% |
| $k_{CL,0}^S$ | 0.68 /day | 80.5% |
| $k_{CL,0}^I$ | 31 /day | 40% |
| $k_{CL,0}^P$ | 0.059 /day | 67% |

**Figure 4:**
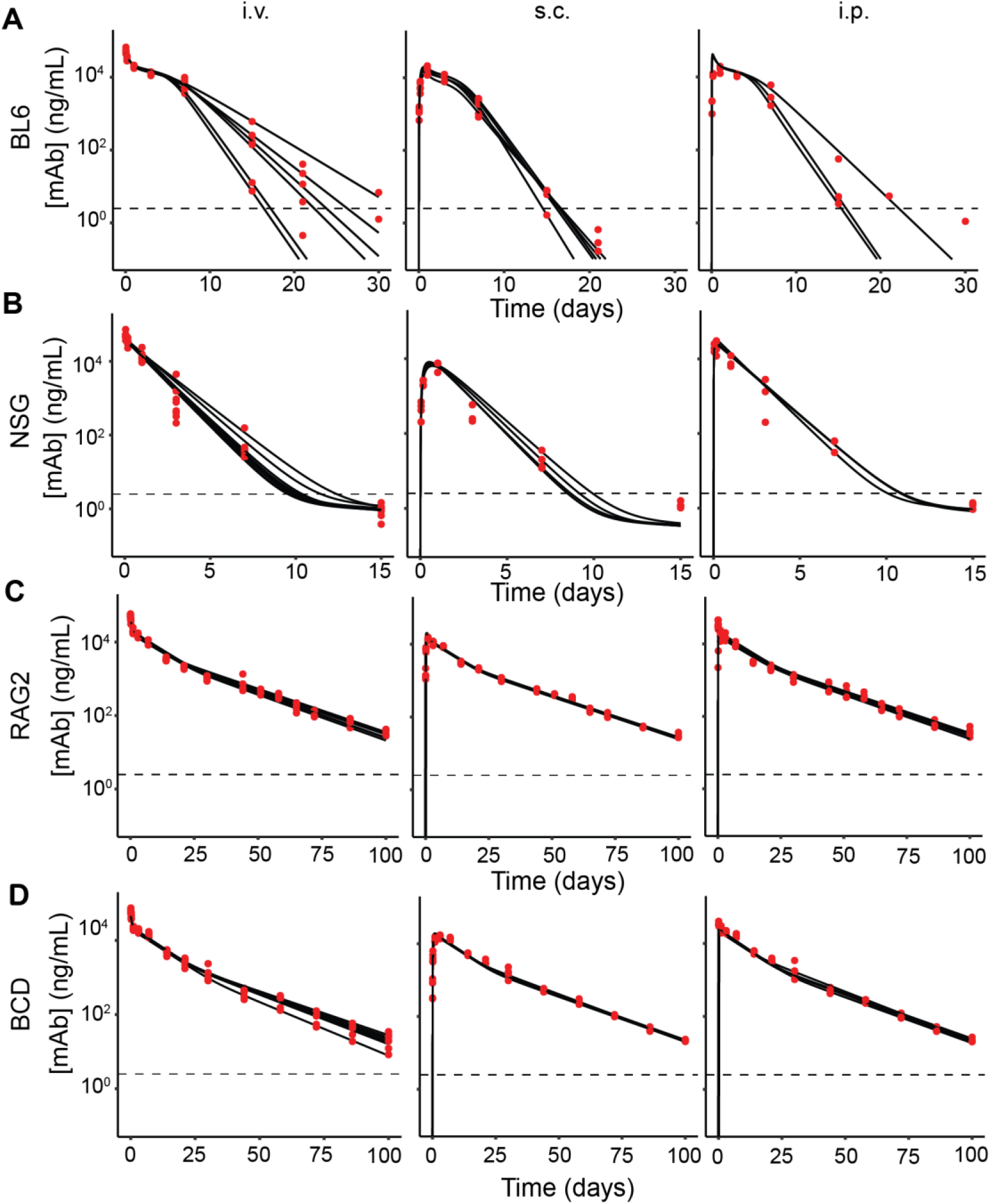
Model fitness to experimental data. A) Fitness to BL6 data. B) Fitness to NSG data. C) Fitness to RAG2 data. D) Fitness to BCD data. Red points: experimental data. Solid black lines: model predictions. Dashed black line: Lower limit of quantification.

Next, we analyzed the estimated parameters and quantified how specific processes in our model change based on each mouse strain. First, the time dependence of clearance in BL6, RAG2, and BCD mice was characterized by calculating the fold change in the clearance rate constant over time based on the estimated parameters for each mouse. In BL6 mice, we predict that the maximum fold-change in clearance varies significantly between mice, with an average increase of around 18x reached between 10 and 20 days after dosing (**Figure 5A**). Conversely, the change in clearance occurs at a much later time in both RAG2 and BCD mice, reaching a maximum decrease around 50 days after dosing. The maximum decrease in clearance varies slightly between RAG2 and BCD mice, at around 70% and 60%, respectively (**Figure 5B**). Next, we used the estimated parameter values for each mouse to quantify the fraction of antibody absorbed into the blood from the peripheral tissue, s.c., and i.p. compartments (**Supplementary Text**). In BL6, RAG2, and BCD mice, close to 100% of antibody that leaves circulation into peripheral tissue is reabsorbed. In NSG mice, however, reabsorption of 3BNC117 into the blood is completely ablated (**Figure 5C**). Interestingly, after s.c. injection, a smaller fraction of 3BNC117 is absorbed in NSG mice compared to other strains, but the effects are far less severe compared to reabsorption after distribution into peripheral tissue (**Figure 5D**). Absorption after i.p. administration, however, is estimated to be quite similar across mouse strains (**Figure 5E**).

**Figure 5:**
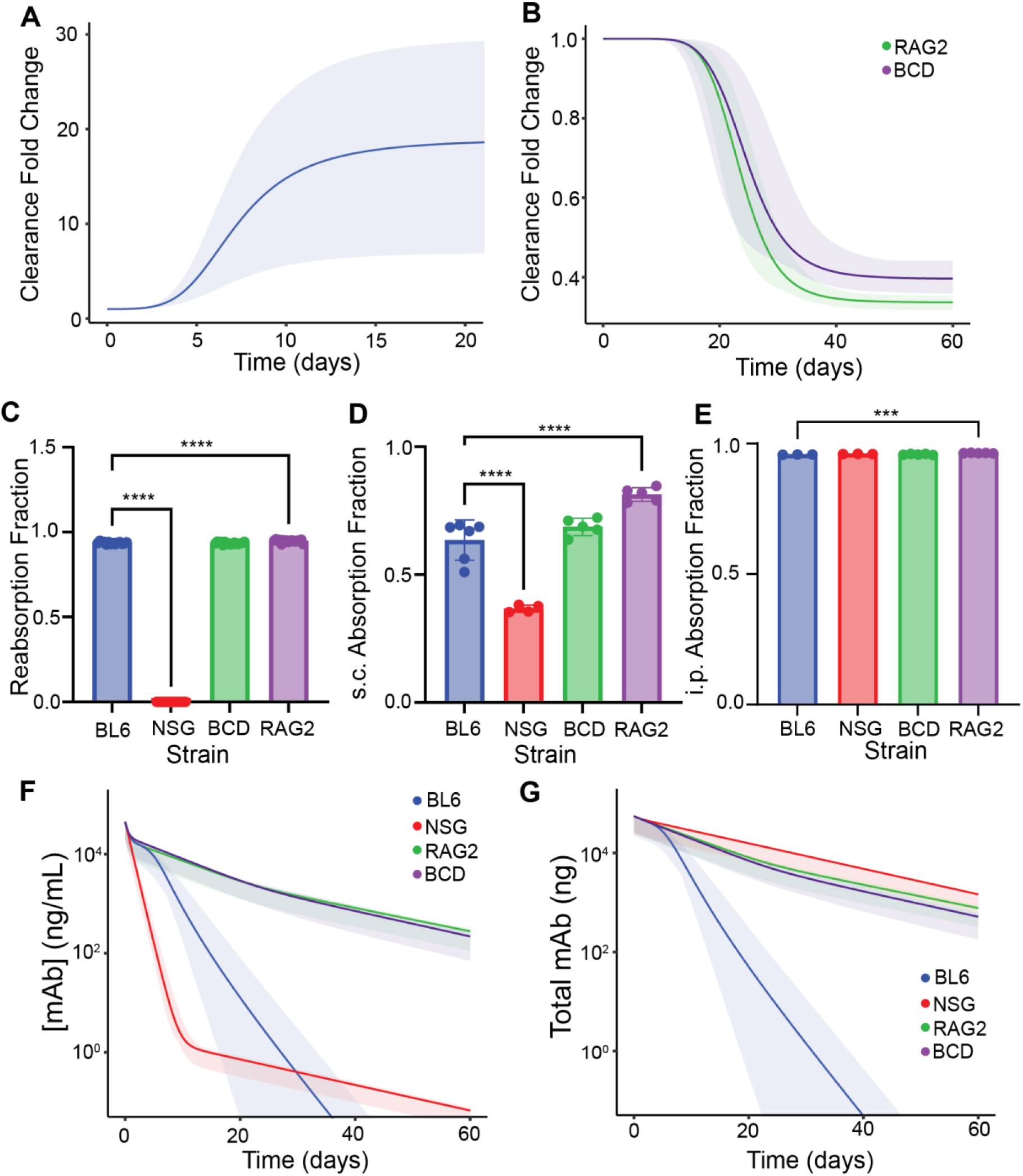
Analysis of optimized model and quantification of strain-dependent differences in PK. A-B) Predicted fold-change in clearance rate constant 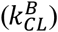 over time in A) BL6 mice and B) RAG2 and BCD mice. C-E) Predicted fractions of 3BNC117 absorbed from C) peripheral tissue compartment, D) s.c. dosing compartment, and E) i.p. dosing compartment. F-G) Comparison of simulated PK profiles for each strain of mouse following bolus i.v. injection. F) Serum 3BNC117 concentrations. G) Total 3BNC117 in body. Solid lines: mean predictions. Shaded regions: 95% interval.

To better understand how the proposed mechanisms in our model influence 3BNC117 disposition in mice, we compared simulated serum mAb concentrations to calculated predictions of total mAb (in ng) in the body over time (**Figure 5F-G**). Simulations reveal that in BL6, RAG2, and BCD mice, serum mAb concentrations closely mirror the predicted amounts of total mAb within the body. The model predicts that increased clearance due to ADA leads to a sharp decrease in 3BNC117 throughout the entire mouse, suggesting that the fast observed clearance in BL6 mice is an elimination-related phenomenon. Conversely, in NSG mice, the sharp decrease in predicted serum mAb is not mirrored by the predicted total amounts of mAb within the body. The fall in serum mAb concentrations is a result of accumulation in peripheral tissues rather than elimination from the body, leading to total mAb remaining high. These results suggest that the apparent decrease in serum titers in NSG mice is a distribution phenomenon and that rapid mAb clearance from the blood does not indicate rapid mAb elimination from the body.

### Simple Allometric Scaling of Model Parameters Suggests NSG Mice are Unpredictive of mAb PK

Next, we used our model to assess the clinical translatability of 3BNC117 in each mouse strain tested by applying simple allometric scaling relationships [24,43] to estimate human model parameters from each mouse strain. Parameters pertaining to compartment volumes were transformed using an allometric scaling exponent of 1, flow rates of 0.85, and rate constants of -0.15 [24,43]. We simulated 30 mg/kg i.v. injections of 3BNC117 in human patients using these estimated parameters (**Figure 6A-B**) and compared the resulting PK profiles to clinical trial data in HIV-negative patients [27]. Clinical trial data was extracted graphically using GraphGrabber2.0.2 (Quintessa). Visual inspection of extracted data reveals that 3BNC117 exhibits a typical mAb PK profile in humans, with an initial rapid distribution phase with a short duration of 1-2 days followed by a slow elimination phase [39,40]. Notably, human predictions using NSG-scaled parameters lack the transition from the rapid distribution phase to the slow elimination phase and deviate significantly from the expected human profile. Human predictions based on RAG2-, BCD-, and BL6-scaled parameters exhibit a rapid distribution phase consistent with human data in terms of both duration and magnitude (**Figure 6A**). The presence of ADA in BL6 mice, however, causes the predicted PK profile to deviate from the expected profile following the onset of ADA formation. Furthermore, the use of NSG-derived parameters predicts a significant accumulation of mAb in peripheral tissues, unlike the other three strains of mice (**Figure 6B**). Altogether, the predicted human PK profiles suggest that NSG mice are an inappropriate mouse model to study mAb PK, whereas BL6 is only appropriate if the effects of ADA on mAb PK are intended to be studied. RAG2 and BCD mice predict a PK profile that is more representative of what is expected.

**Figure 6:**
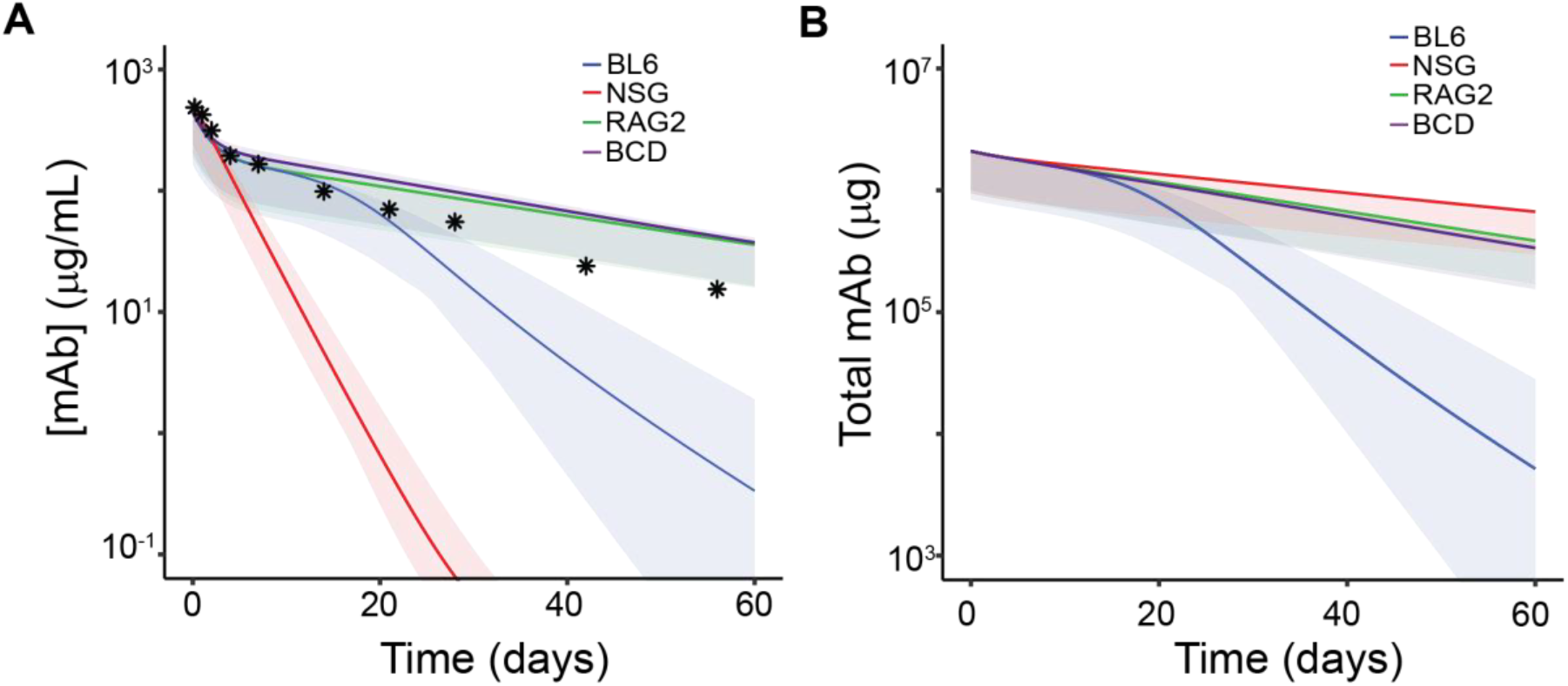
Simulated 3BNC117 PK following bolus i.v. injection in humans estimated with simple allometric scaling of model parameters. A) Predictions of serum 3BNC117 concentrations v. time. B) Predictions of total 3BNC117 within the body v. time. Solid lines: mean predicted values. Shaded regions: 95% interval. Stars: Extracted PK data from clinical trial [27].

## Discussion

In this work, we generated a detailed dataset of 3BNC117 PK after bolus i.v., s.c., and i.p. injection in four strains of mice to investigate the role of immune system composition in mAb PK. With a specific emphasis on providing a quantitative comparison of how certain processes in each mouse strain vary, we developed an mPBPK model and applied nonlinear mixed effects modeling with strain-based covariates to estimate the differences in model parameters between strains. Our model incorporated time-dependent clearance due to the suspected onset of ADA in BL6 mice and impaired reabsorption of mAb into the blood in NSG mice, leading to accumulation in peripheral tissues. By estimating model parameters specific to each mouse strain and applying simple allometric scaling methods, we estimated the corresponding human PK profile of 3BNC117 based on each mouse strain. Comparing the predicted profiles to clinical trial data in HIV-negative patients, we ultimately found that the fully immunocompromised NSG mice exhibited mAb PK that leads to significantly different predicted human PK. The results, therefore, suggest that preclinical investigation of mAb PK should not be conducted in NSG mice.

The abnormal PK of mAbs in NSG mice has been previously illustrated [25,26], and our results complement these findings by presenting mAb PK following other routes of administration and characterizing the PK profiles with mathematical modeling. Sharma et al. [25] previously demonstrated through the use of in vivo imaging that several cancer-targeted mAbs accumulate in the spleen, liver, and bone marrow in NOD SCID and NSG mice in a Fc-binding dependent manner. In a similar study, Li et al. [26] evaluated the PK of several mAbs and antibody-drug conjugates and found that NSG mice exhibited significantly faster clearance than other strains. Interestingly, preventing FcγR binding between the biologic and NSG mice cells led to normal PK profiles. Altogether, these studies suggest that mAbs are sequestered in specific peripheral tissues in NSG mice due to high levels of Fc region binding, which is responsible for the observed differences in PK.

In our work, we expanded on these findings by i) investigating the influence of mouse strain on the PK of mAbs following alternative routes of administration, i.e., s.c. or i.p. bolus injections, and ii) quantifying how specific processes during mAb absorption, distribution, and elimination change based on mouse strain. Somewhat unexpectedly, estimated model parameters predict that only reabsorption of mAb after distribution into peripheral tissues from the blood is impaired in NSG mice. We suspect this is because FcγR-dependent mAb sequestration occurs in specific tissues such as the spleen, liver, and bone marrow, and the mAb is only exposed to these areas during distribution from the blood into peripheral tissues rather than absorption following e.v. bolus injections. In addition to this, the fluid administered when dosing into e.v. compartments may also facilitate absorption into the lymphatic system, further preventing accumulation within the injected tissue.

Preclinical characterization of mAbs or other biologics is often evaluated in immunocompromised mouse models to control for xenogeneic responses to the therapeutic being delivered. During these experiments, selecting a mouse model that will exhibit mAb PK that is predictive of what is expected in clinical translation is critical. By estimating strain-specific PK parameters with our mPBPK model, we approximated the translatability of each mouse model through allometric scaling to predict human model parameters based on each strain. We selected 3BNC117 for this study because the variable region of the antibody does not bind any targets in mice, and clinical trial data is available in HIV-patients such that target binding is also absent [27]. Therefore, dosing of 3BNC117 in both mice and HIV-humans results in PK profiles unaffected by mAb target binding, or target-mediated drug disposition (TMDD) [44]. Allometrically scaled model predictions demonstrated that NSG mice exhibit mAb PK that is not predictive of human dosing. Predicted human 3BNC117 time courses leveraging data collected from NSG mice were far different from clinical trial data [27]. BL6-, RAG2-, and BCD-derived predictions were all consistent with human PK at early time points, but the presence of ADA caused BL6 predictions to deviate. Overall, our investigation into the strain dependence of mAb PK via mathematical modeling allowed us to evaluate the translatability of each mouse strain quantitatively. Importantly, our results demonstrate that RAG2 and BCD mice were the most suitable mouse strains for preclinical mAb characterization and emphasize the importance of mouse strain selection in preclinical experiments.

Our findings are particularly important regarding the preclinical characterization of drug delivery systems for the controlled release of mAbs and other large biologics, including drug delivery devices and gene therapies. The development of these technologies is of great interest today because of their ability to maintain therapeutic drug concentrations at target tissues and safe concentrations in circulation. Accordingly, these technologies are in development for the treatment of a variety of diseases including cancer [15,16,45–47], macular degeneration [17,48,49], inflammatory disease [49,50], and HIV [51,52]. In all these cases, preclinical characterization of the delivered therapeutic’s PK is essential. Immunocompromised mouse models are often selected for these experiments to allow study of therapeutic PK in the absence of any xenogeneic responses against the drug or device itself. However, selecting a small animal model that exhibits translatable PK is critical to streamline preclinical experimentation. An ideal KO model for preclinical assessment of PK is sufficiently immunocompromised to mitigate the ADA or FBR responses but still exhibits clinically translatable PK. By characterizing the effects of immune KO on 3BNC117 in detail, we emphasize the importance of selecting an appropriate mouse model and provide strong evidence that the NSG mouse model is inappropriate for PK testing. We also demonstrate that RAG2 and BCD mice are well suited for preclinical experimentation of those tested in this work.

## Author Contributions

Jonathon DeBonis: Conceptualization, Methodology, Software, Formal Analysis, Writing – Original Draft, Visualization. Anthony Davis: Conceptualization, Methodology, Validation, Formal Analysis, Investigation, Writing – Original Draft, Visualization. Zeshi Wang: Investigation. Cody Fell: Conceptualization, Writing – Review & Editing. Michael Diehl: Conceptualization, Methodology, Writing – Review & Editing, Funding acquisition. Oleg Igoshin: Conceptualization, Methodology, Writing – Review & Editing, Funding acquisition. Omid Veiseh: Conceptualization, Methodology, Writing – Review & Editing, Funding acquisition.

## Acknowledgements

This work was supported by the Gates Foundation (INV-051204) and by Welch Foundation Award C-1995 (to OAI). Parameter estimation was performed using the Big-Data Private-Cloud Research Cyberinfrastructure MRI-award funded by NSF under grant CNS-1338099 and by Rice University’s Center for Research Computing (CRC). This material is based upon work supported by the National Science Foundation Graduate Research Fellowship Program awarded to Anthony Davis. Any opinions, findings, and conclusions or recommendations expressed in this material are those of the authors and do not necessarily reflect the views of the National Science Foundation. The 3BNC117 protein was provided by the CAVD Protein Production Central Services Facility at the Duke Human Vaccine Institute (DHVI). The CAVD Protein Production Facility is supported by a Collaboration for AIDS Vaccine Discovery grant (INV-045468) from the Gates Foundation.

## Supplementary Information

Supplementary information to this article can be found in the attached PDF file “Supplementary Information.pdf” and the attached excel file “Supplementary Data.xlsx”.

## SUPPLEMENTARY INFORMATION

### Supplementary Tables

**Table S1:**
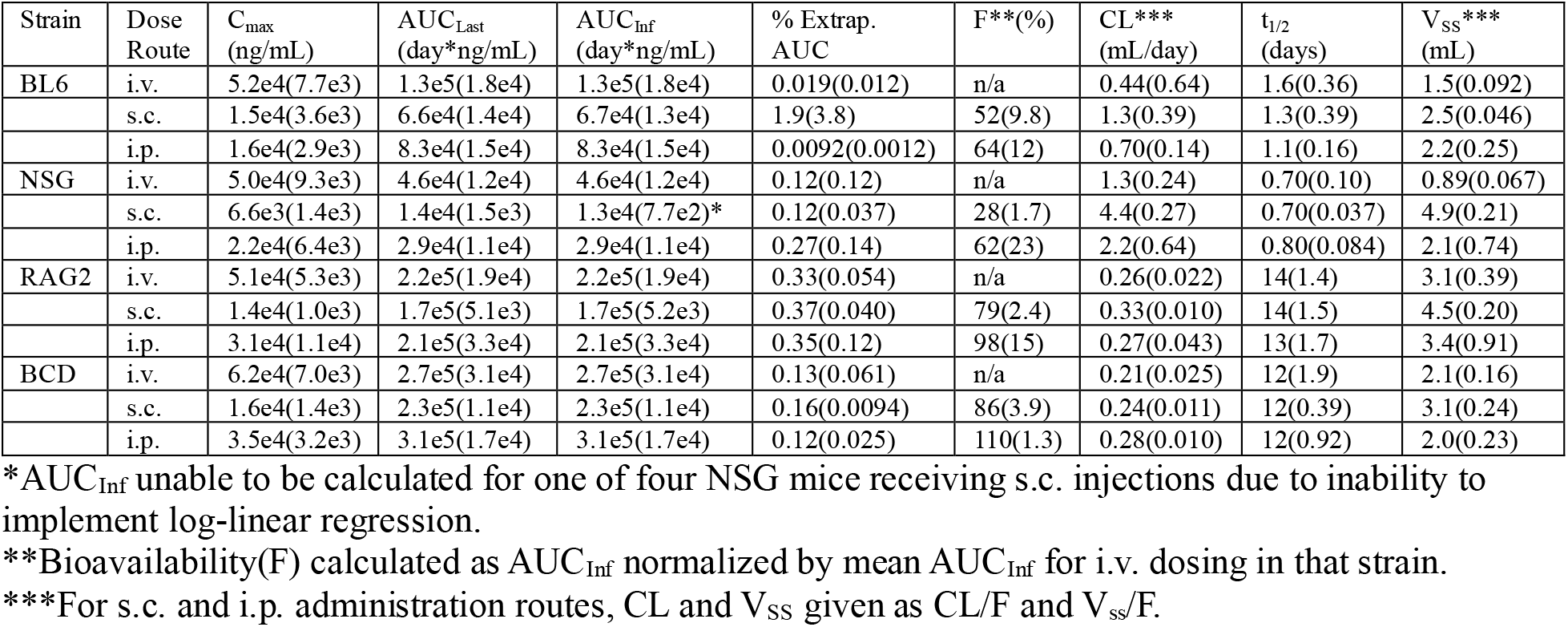
NCA Parameters & Statistics Based on Strain of Mouse. Values reported to two significant digits. () indicates standard deviation. C_max_ = maximum measured concentration. AUC_Last_ = area under the concentration v. time curve up to last measured concentration. AUC_Inf_ = extrapolated AUC to time infinity. % Extrap. AUC = percentage of AUC_Inf_ extrapolated using log-linear regression. F = bioavailability. CL = clearance. t_1/2_ = terminal half life. V_ss_ = steady-state volume of distribution.

| Strain | Dose Route | $C_{\max}$<br>(ng/mL) | $AUC_{\text{Last}}$<br>(day*ng/mL) | $AUC_{\text{Inf}}$<br>(day*ng/mL) | % Extrapol.<br>AUC | F** (%) | CL***<br>(mL/day) | $t_{1/2}$<br>(days) | $V_{\text{ss}}$***<br>(mL) |
| --- | --- | --- | --- | --- | --- | --- | --- | --- | --- |
| BL6 | i.v. | 5.2e4(7.7e3) | 1.3e5(1.8e4) | 1.3e5(1.8e4) | 0.019(0.012) | n/a | 0.44(0.64) | 1.6(0.36) | 1.5(0.092) |
|  | s.c. | 1.5e4(3.6e3) | 6.6e4(1.4e4) | 6.7e4(1.3e4) | 1.9(3.8) | 52(9.8) | 1.3(0.39) | 1.3(0.39) | 2.5(0.046) |
|  | i.p. | 1.6e4(2.9e3) | 8.3e4(1.5e4) | 8.3e4(1.5e4) | 0.0092(0.0012) | 64(12) | 0.70(0.14) | 1.1(0.16) | 2.2(0.25) |
| NSG | i.v. | 5.0e4(9.3e3) | 4.6e4(1.2e4) | 4.6e4(1.2e4) | 0.12(0.12) | n/a | 1.3(0.24) | 0.70(0.10) | 0.89(0.067) |
|  | s.c. | 6.6e3(1.4e3) | 1.4e4(1.5e3) | 1.3e4(7.7e2)* | 0.12(0.037) | 28(1.7) | 4.4(0.27) | 0.70(0.037) | 4.9(0.21) |
|  | i.p. | 2.2e4(6.4e3) | 2.9e4(1.1e4) | 2.9e4(1.1e4) | 0.27(0.14) | 62(23) | 2.2(0.64) | 0.80(0.084) | 2.1(0.74) |
| RAG2 | i.v. | 5.1e4(5.3e3) | 2.2e5(1.9e4) | 2.2e5(1.9e4) | 0.33(0.054) | n/a | 0.26(0.022) | 14(1.4) | 3.1(0.39) |
|  | s.c. | 1.4e4(1.0e3) | 1.7e5(5.1e3) | 1.7e5(5.2e3) | 0.37(0.040) | 79(2.4) | 0.33(0.010) | 14(1.5) | 4.5(0.20) |
|  | i.p. | 3.1e4(1.1e4) | 2.1e5(3.3e4) | 2.1e5(3.3e4) | 0.35(0.12) | 98(15) | 0.27(0.043) | 13(1.7) | 3.4(0.91) |
| BCD | i.v. | 6.2e4(7.0e3) | 2.7e5(3.1e4) | 2.7e5(3.1e4) | 0.13(0.061) | n/a | 0.21(0.025) | 12(1.9) | 2.1(0.16) |
|  | s.c. | 1.6e4(1.4e3) | 2.3e5(1.1e4) | 2.3e5(1.1e4) | 0.16(0.0094) | 86(3.9) | 0.24(0.011) | 12(0.39) | 3.1(0.24) |
|  | i.p. | 3.5e4(3.2e3) | 3.1e5(1.7e4) | 3.1e5(1.7e4) | 0.12(0.025) | 110(1.3) | 0.28(0.010) | 12(0.92) | 2.0(0.23) |

\* $AUC_{\text{Inf}}$ unable to be calculated for one of four NSG mice receiving s.c. injections due to inability to implement log-linear regression.
\*\*Bioavailability(F) calculated as $AUC_{\text{Inf}}$ normalized by mean $AUC_{\text{Inf}}$ for i.v. dosing in that strain.
\*\*\*For s.c. and i.p. administration routes, CL and $V_{\text{ss}}$ given as CL/F and $V_{\text{ss}}$ /F.

**Table S2:** Description of mPBPK model parameters including initial guesses for estimated parameters. Values are reported to two significant digits.

| Parameter | Meaning | Value/Initial Guess | Source |
| --- | --- | --- | --- |
| $j_{TB}$ | Fluid flow rate from blood to peripheral tissue (mL/day) | 3.0 | Obtained from naïve pooled fitting. |
| $j_{PT}$ | Fluid flow from peripheral tissue to peripheral lymphatics (mL/day) | 0.15 | Obtained from naïve pooled fitting. |
| $\delta_{NSG}$ | Scaling factor for $j_{PT}$ in NSG mice. | 0.0010 | Obtained from naïve pooled fitting. |
| $j_{PS}$ | Fluid flow rate from s.c. dosing compartment to peripheral lymphatics (mL/day) | 1.0 | Obtained from naïve pooled fitting. |
| $j_{PI}$ | Fluid flow rate from i.p. dosing compartment to peripheral lymphatics (mL/day) | 10 | Obtained from naïve pooled fitting. |
| $j_{BL}$ | Fluid flow rate from central lymphatic compartment to blood (mL/day) | 2.4 (fixed) | Taken from Trevaskis et al. [1] |
| $j_{LP}$ | Fluid flow rate from peripheral lymphatic compartment to central lymphatic compartment (mL/day) | 2.4 (fixed) | Assumed to be equal to $j_{BL}$. |
| $k_{CL,0}^S$ | First-order elimination rate constant from s.c. dosing compartment at median mouse weight(1/day) | 1.0 | Obtained from naïve pooled fitting. |
| $k_{CL,0}^I$ | First-order elimination rate constant from i.p. dosing compartment at median mouse weight (1/day) | 10.0 | Obtained from naïve pooled fitting |
| $k_{CL,0}^P$ | First-order elimination rate constant from peripheral tissues, peripheral lymphatic, and central lymphatic compartments at median mouse weight (1/day) | 0.029 | Obtained from naïve pooled fitting. |
| $k_{CL,0}^B$ | First order elimination rate constant from central/blood compartment at median mouse weight (1/day) | 0.32 | Obtained from naïve pooled fitting. |
| $T_{max}^{R/B}$ | Parameter governing maximum fold-change in clearance rate constant over time in RAG2 and BCD mice (unitless) | -1.0 | Obtained from naïve pooled fitting. |
| $T50^{R/B}$ | Parameter governing time at which clearance reaches 50% of maximum fold change in RAG2 and BCD mice (days) | 21 | Obtained from naïve pooled fitting |
| $g^{R/B}$ | Hill coefficient for clearance fold-change equation in RAG2 and BCD mice (unitless) | 3.2 | Obtained from naïve pooled fitting |
| $T_{max}^{BL6}$ | Parameter governing maximum fold-change in clearance rate constant over time in BL6 mice (unitless) | 1.0 | Obtained from naïve pooled fitting. |
| $T50^{BL6}$ | Parameter governing time at which clearance reaches 50% of maximum fold change in BL6 mice (days) | 6.0 | Obtained from naïve pooled fitting |
| $g^{BL6}$ | Hill coefficient for clearance fold-change equation in BL6 mice (unitless) | 3.9 | Obtained from naïve pooled fitting |
| $V_{B,0}$ | Volume of distribution of central/blood compartment at median mouse weight (mL) | 0.96 | Obtained from naïve pooled fitting |
| $V_T$ | Volume of peripheral tissue compartment (mL) | 0.050 | Approximated |
| $V_P$ | Volume of peripheral lymphatic compartment (mL) | 0.050 | Approximated |
| $V_L$ | Volume of central lymphatic compartment (mL) | 0.050 | Approximated |
| $V_S$ | Volume of s.c. dosing compartment (mL) | 0.10 | Set to volume of fluid injected |
| $V_I$ | Volume of i.p. dosing compartment. | 0.10 | Set to volume of fluid injected |
| $BW$ | Body weight of individual mouse being simulated (g) | variable | Determined by experimental measurements |
| $BW_{median}$ | Median body weight of all mice in experiments (g) | 19 | Determined by experimental measurements |

### Supplementary Figures

**Figure S1:**
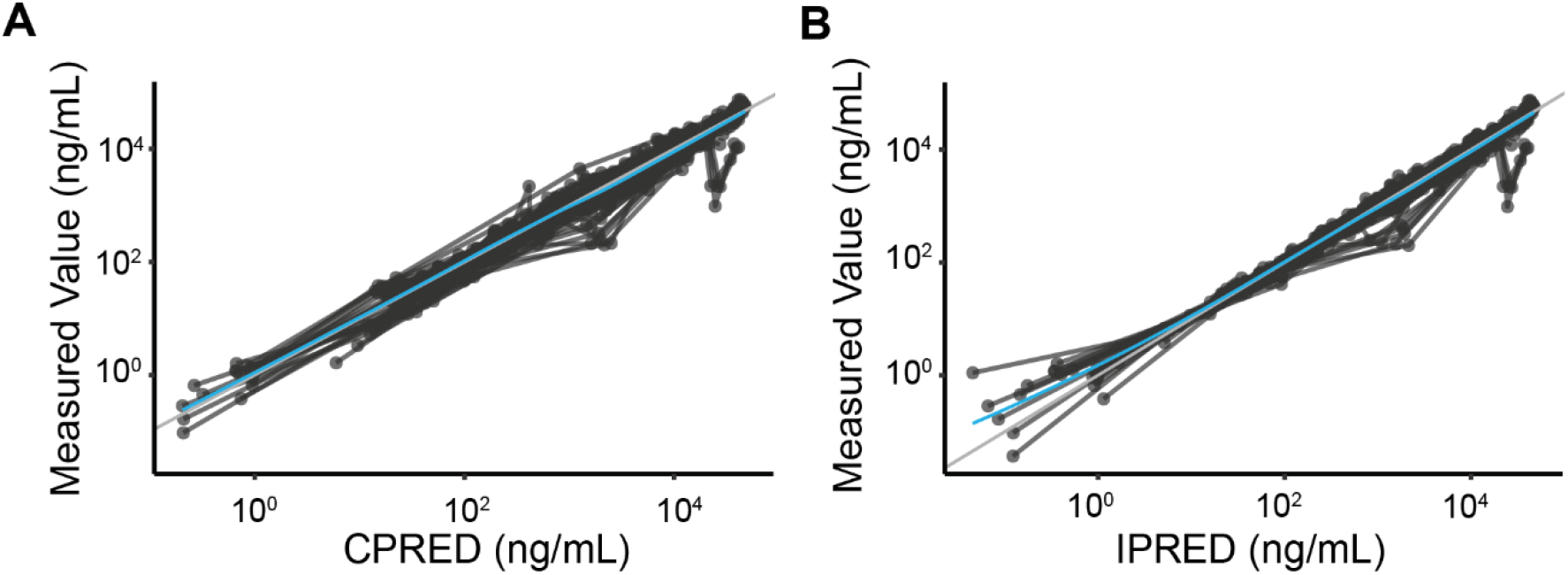
Measured serum concentrations v. model predicted values. A) Experimental concentrations v. predicted values using population parameters. B) Experimental concentrations v. predicted values using individual mouse parameters. CPRED: Population predicted values. IPRED: Individual predicted values.

**Figure S2:**
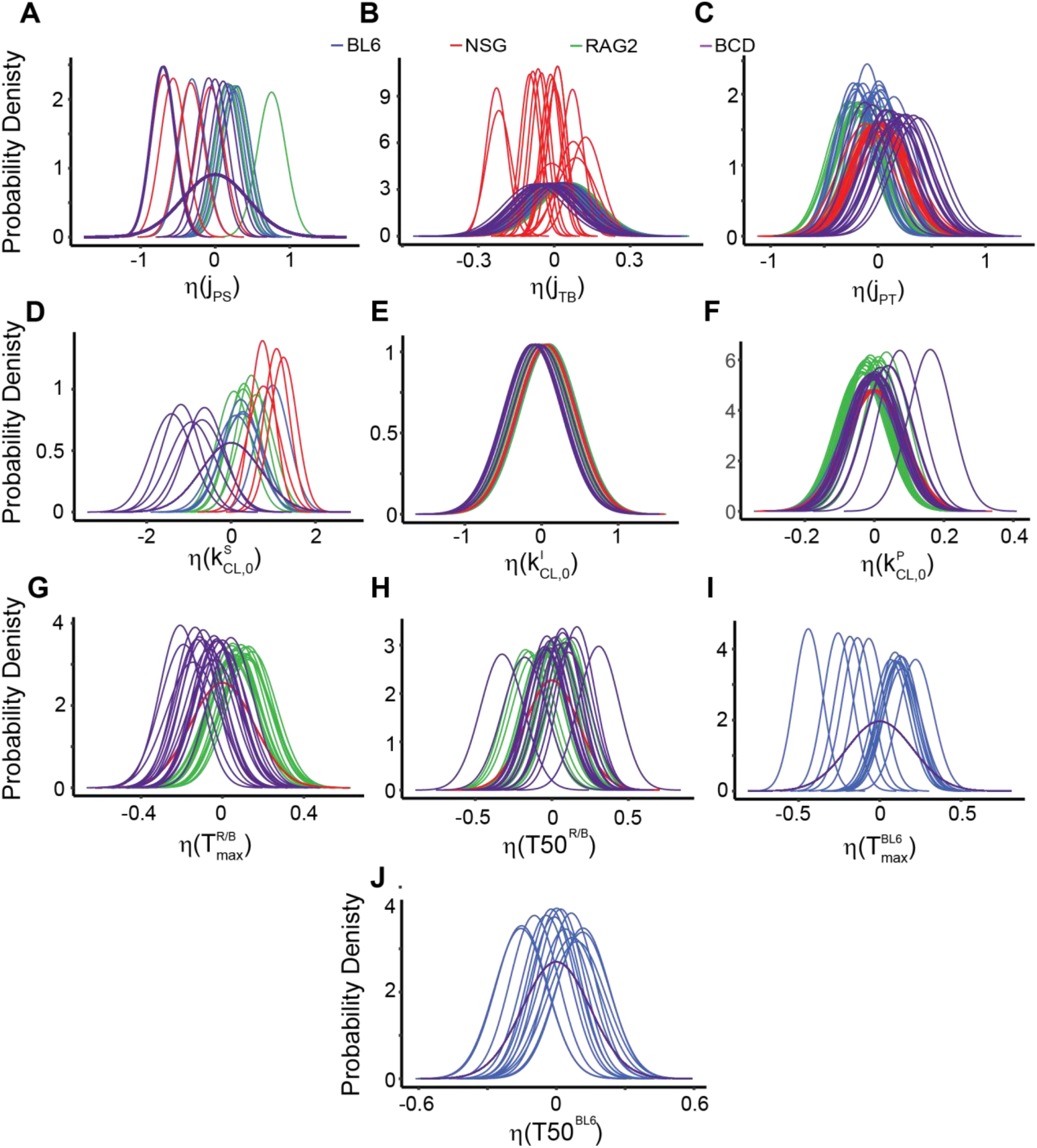
Empirical Bayes Estimates (EBE) of individual random effect model parameters. *η*(*p*) indicates, for a given parameter 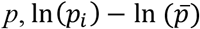, where *p*_*i*_ is the individual estimated value of *p* and 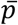 is the population estimate of parameter *p*. EBEs presented for all random effect parameters: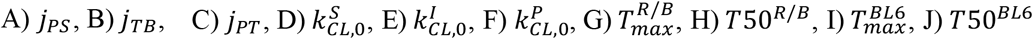. Blue curves: BL6 mice, red curves: NSG mice, green curves: RAG2 mice, purple curves: BCD mice.

### Supplementary Text

#### Derivation of Model Equations

The minimal physiologically based pharmacokinetic (mPBPK) model developed in this work consists of six ordinary differential equations governing transport and elimination between compartments. The model contains six compartments: The model consists of six total compartments: the central/blood compartment (B), the peripheral tissue compartment (T), the peripheral lymphatic compartment (P), the central lymphatic compartment (L), the subcutaneous dosing compartment (S), and the intraperitoneal dosing compartment (I). In the sections below, we derive and explain the terms used for transport and elimination in our model.

##### Dosing

Dosing in our model was represented by setting the initial concentration in the dosing compartment equal to the amount of mAb administered divided by the volume of the corresponding compartment according to:

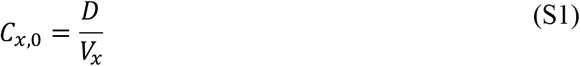

Where *C*_*x*_ is the concentration of antibody in compartment *x* (ng/mL) and *V*_*x*_ is the volume of compartment *x* (mL).

##### Transport Between Compartments

In our model, transport between compartments is represented by unidirectional fluid flow between compartments. In the case where fluid flows from compartment *i* to compartment *j*, the concentration of mAb in each compartment is described by:

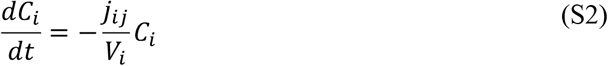

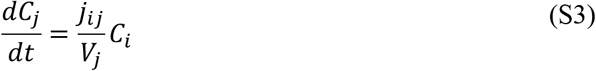

Where *j*_*ij*_ is the rate of fluid flow from compartment *i* to compartment *j* (mL/day). The topology of fluid flow between compartments was based on models in the literature [2,3].

##### Elimination from Compartments

Because mAbs are primarily eliminated from the body through cellular uptake and lysosomal degradation, elimination terms are included for all compartments in our model. Elimination is described as a first order process according to:

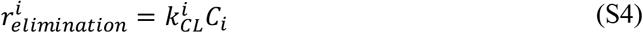

Where 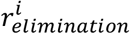 is the rate of elimination (ng/mL/day) from compartment *i*, and 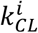 is the first order rate constant describing elimination from compartment *i*. We assume that each compartment is described by a separate elimination rate constant, except for the peripheral tissue (T), peripheral lymphatic (P), and central lymphatic (L) compartments, which share the same elimination rate constant 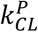. Furthermore, due to the observed time-dependence of mAb clearance observed in experimental data, clearance from the central/blood compartment (B) for C57BL6, RAG2, and BCD mice was given time dependence described by:

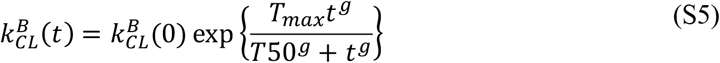

Where the parameter *T*_*max*_ determines the maximum fold-change in the clearance rate constant (unitless), *T*50 is the time at which clearance reaches 50% of its maximum change (days), *g* is a hill coefficient, and 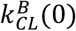 is the value of the clearance rate constant at time zero (1/day). This equation was based on previous models in the literature describing time-dependent mAb elimination [4,5]

##### Model Covariates

Our model incorporates mouse starting weight and mouse strain as covariates. Mouse weight influences the values of clearance rate constants and the volume of the central compartment according to:

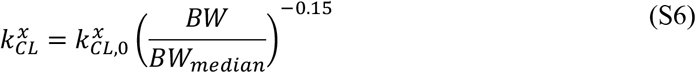

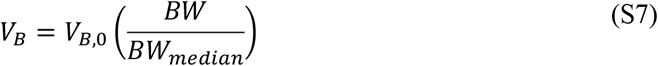

Where *BW* is the mouse’s body weight (g) and *BW*_*median*_ is the median body weight of the mice being studied (g). These equations and the selected allometric scaling exponents were based on literature values [6,7].

In addition to mouse body weight as a covariate, the mouse strain was included as a covariate and influenced the fluid flow rate from the peripheral tissue compartment to the peripheral lymphatic compartment (*j*_*PT*_) and the parameters governing clearance time-dependence (*T*_*max*_, *T*50, *g*). Impaired ability of NSG mice to reabsorb mAb from peripheral tissue is represented by a scaling of the fluid flow rate *j*_*PT*_ by the factor *δ*_*NSG*_. Time-dependence was only observed in BL6, RAG2, and BCD PK data; thus, time dependence was only included in the model for these three strains. In BL6 mice clearance appeared to increase around one week after dosing, whereas in RAG2 and BCD mice clearance appeared to decrease around 40 days after dosing. Accordingly, the clearance kinetics in BL6 mice were described by 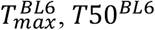, and *g*^*BL*6^ and in RAG2/BCD mice were described by 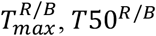, and *g*^*R/B*^.

#### Derivation of Absorption Fractions

Estimated model parameters in the main text are used to calculate the predicted fraction of antibody that is absorbed into the blood from the peripheral tissue, subcutaneous, and intraperitoneal compartments. Consider a single compartment *x* with concentration of mAb *C*_*x*_ and volume *V*_*x*_ where mAb leaves the compartment via two routes: i) transport to compartment *y* mediated by fluid flow rate *j*_*yx*_ and ii) degradation governed by the first-order degradation rate constant 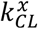. The fraction of mAb that reaches compartment *y* is equivalent to the flux of mAb to compartment *y* to the total flux leaving the compartment. This quantity (*f*_*x*_) is given by:

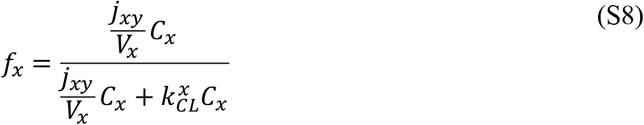

This equation in its simplified form is:

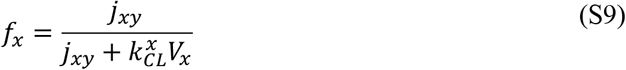

Importantly, this quantity is not dependent on the current concentration of mAb in compartment *x* and is solely a function of the compartment volume, the fluid flow rate to the next compartment, and the elimination rate constant. The fraction of antibody that reaches the blood from a given compartment in our model is equivalent to the product of all *f*_*x*_ values for each compartment the mAb is exposed to before reaching the blood. From the peripheral tissue, mAb is exposed to the peripheral tissue compartment, the peripheral lymphatic compartment, and the central lymphatic compartment. Therefore, the fraction of mAb reabsorbed into the blood from the peripheral tissue compartment (*f*_*reabsorbed*_) is given by:

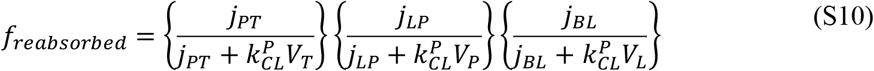

Similarly, the fractions of mAb absorbed following s.c. (*f*_*s*.*c*._) or i.p. (*f*_*i*.*p*._) bolus mAb injection are given by:

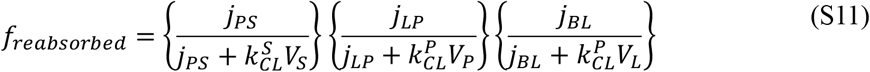

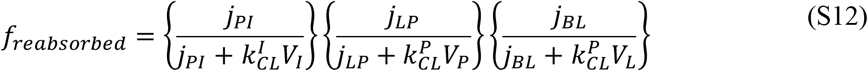

